# Timing, movement, and reward contributions to prefrontal and striatal ramping activity

**DOI:** 10.1101/2025.07.02.662841

**Authors:** Alexandra Bova, Matthew A. Weber, Mackenzie M. Spicer, Vaibhav Khandelwal, Rachael A. Volkman, Madison McMurrin, Julia Kryca, Kartik Sivakumar, Braedon Q. Kirkpatrick, Qiang Zhang, Nandakumar S. Narayanan

**Author notes:** **Corresponding Author**: Dr. Nandakumar Narayanan, MD, PhD, 169 Newton Road, Pappajohn Biomedical Discovery Building—5336 University of Iowa, Iowa City, IA 52245. **Conflict of Interest**: There are no conflicts of interest.

## Abstract

Across species, prefrontal and striatal neurons exhibit time-dependent ramping activity, defined as a consistent monotonic change in firing rate across temporal intervals. However, it is unclear if ramping activity is related to the cognitive process of estimating time, or to other behavioral factors such as anticipating reward or regulating movements. Here, we harnessed two novel approaches to determine how these factors contribute to prefrontal and striatal ramping activity in mice performing an interval timing task. First, to determine how movement contributes to ramping activity, we tracked movement velocity using DeepLabCut as well as task-specific movements while recording prefrontal or striatal ensembles during interval timing. We found that time was more accurately decoded by ramping neurons than movement-modulated neurons, with the exception of prefrontal velocity-modulated neurons. Second, to disambiguate temporal signals from anticipatory reward signals we compared activity patterns in neurons that were recorded during interval timing to the same neurons recorded during a Pavlovian conditioning task. We found more ramping activity and more accurate temporal decoding by neuronal ensembles during interval timing compared to Pavlovian conditioning. Together, these data quantify contributions of time estimation, movement, and reward anticipation in prefrontal and striatal ensembles, and they suggest that ramping is a cognitive signal that estimates time. Our results provide insight into how prefrontal and striatal ensembles multiplex information to effect temporal control of action.

## INTRODUCTION

Mammalian behavioral repertoires unfold over time, during which neural systems must organize movements and anticipate consequences. During behavior that occurs over a temporal interval, neurons in the mammalian prefrontal cortex and the striatum exhibit time-dependent ramping activity, defined as a consistent monotonic increase or decrease in the firing rate across a temporal interval^1,2^. Ramping activity is observed in humans, primates, and rodents^2–5^. Data-driven techniques such as principal component analysis (PCA) frequently find that ramping activity explains the largest prefrontal and striatal variance during timed behaviors^5–8^. Because prefrontal and striatal neurons are required for motivated behavior and can malfunction in human neurological and psychiatric diseases, elucidating prefrontal and striatal neuronal activity is critical to help us better understand human disease, develop new neurophysiological markers, and inspire next-generation therapies.

The behavioral significance of time-dependent ramping activity is unclear. One possibility is that ramping activity represents the passage of time as it scales with temporal intervals, as predicted by drift-diffusion models accounting for accumulating temporal evidence^5,9,10^. A second possibility is that ramping activity simply reflects movement, because motor sequences are organized in time^11^. A third possibility is that ramping activity does not reflect ongoing actions, but instead encodes the anticipation of a future reward^12^. Resolving these possibilities is difficult: until recently, movement has rarely been tracked during behavior with neuronal recordings, and it is challenging to dissociate reward anticipation from timing- and movement-related neuronal signals because most behavioral paradigms are motivated by reward.

We directly addressed these issues by recording prefrontal and striatal neurons as mice performed an interval timing task in which they estimate an interval of several seconds by making a motor response. We advance previous corticostriatal recordings in two ways: 1) to dissociate ramping activity from movement, we tracked movement using DeepLabCut^13,14^, and 2) to dissociate ramping from anticipatory reward activity, we recorded neuronal activity in mice trained to perform not only an operant interval timing task but also a Pavlovian conditioning task, in which cues signaled an upcoming reward delivery. We recorded neuronal activity in the mouse prefrontal cortex (PFC) and dorsomedial striatum (DMS) during interval timing and found prominent time-dependent ramping activity, as well as velocity- and nosepoke-related activity. Crucially, we found that ramping neurons decoded time more accurately than movement-modulated neurons, with the exception of velocity-modulated neurons in the PFC. When recording neuronal activity during the Pavlovian conditioning task, we observed fewer ramping neurons than during interval timing. Furthermore, we found that prefrontal and striatal ensembles decoded time more accurately during interval timing than during the Pavlovian conditioning task. These data establish time-dependent ramping as a key temporal code.

## METHODS

### Mice

All procedures were approved by the Institutional Animal Care and Use Committee (IACUC) at the University of Iowa (#3052039). We used two cohorts of mice: 1) 8 mice (4 female) for recordings from the prefrontal cortex (PFC; 7 wild-type C57BL6/J and 1 D1-IRES-Cre mice; different recording sessions from these mice were included in^40^); and 2) 11 mice (5 female) for recordings from dorsomedial striatum (DMS; 7 wild-type C57BL6/J and 4 DAT-IRES-Cre mice). Mice were food restricted to 85% of their starting bodyweight, had access to *ad libitum* water, and were housed individually post-surgery, according to methods described in detail previously^15,16^.

### Interval timing switch task

We used a mouse-optimized interval timing task described in detail previously^16–18^. Mice were trained in sound-attenuating chambers, with two nosepokes on either side of a central food magazine on the front wall, and a third nosepoke at the center of the back wall. All nosepokes contained lights and a cue light was positioned above each of the two front nosepokes (Fig. 1A; MedAssociates, St. Albans, VT). The nosepokes and the food magazine contained infrared beams to register mouse entries and exits. Training began with fixed-ratio nosepoke response trials to earn rewards (20 mg sucrose pellets; Bio-Serv, Flemington, NJ).

After fixed-ratio response trials, mice began training in the interval timing switch task. Trial availability was signaled by illumination of the back nosepoke. Mice initiated trials by responding at the back nosepoke which triggered the cue lights, front nosepoke lights, and an auditory tone (3kHz, 72 dB). Cues were identical on all trials and were sustained for the entire trial duration. On 50% of randomly assigned trials, cues remained on for 6 seconds, and the first nosepoke at the designated “short” nosepoke (counterbalanced left or right across mice) after 6 seconds was rewarded. On the other 50% of trials, cues remained on for 18 seconds, and mice initially responded at the short nosepoke and then *switched* to the “long” nosepoke. A nosepoke at the long nosepoke after 18 seconds earned a reward. This type of trial is defined as a *switch trial*, and the time at which mice exit the short nosepoke before responding at the long nosepoke is defined as the switch *response time*. Because cues do not differ between short and switch trials except for their duration, switch response times are guided by the mouse’s internal estimate of time and reflect temporal control of action (Fig. 1A–B).

Only switch trials were analyzed. The *coefficient of variation (CV)* was calculated as the standard deviation of switch response times divided by the mean switch response time. The *number of nosepokes* at the short and long nosepokes were only counted for switch trials (between 0–18 seconds). The *percent of correct switch trials* was calculated as the number of switch trials in which mice received a reward divided by the total number of switch trials in a session.

### Pavlovian conditioning task

Following interval timing recordings, the same mice were trained in a Pavlovian conditioning task. Pavlovian conditioning sessions took place in the same chamber as the interval timing task, but the context was changed by fitting 3D-printed walls into the chamber to cover the front and back nosepokes. During the 30-minute Pavlovian conditioning task sessions, mice were randomly presented concurrent light and auditory cue tones for 7 seconds, followed by delivery of a sucrose pellet reward. Notably, the auditory cue tone was distinct from the cue tone used during interval timing sessions (6 kHz, 72 dB). *Latency to retrieve reward* was measured as the time between reward delivery and entry into the food magazine. Mice were trained for an average of 5.0 ± 0.7 Pavlovian conditioning sessions before recordings took place (PFC: 6.4 ± 1.9 sessions, DMS: 4.1 ± 0.3 sessions), with an average of 19.9 ± 3.4 calendar days between interval timing sessions and Pavlovian recordings (PFC: 16.4 ± 3.1, DMS: 22.5 ± 5.4 days).

### Movement tracking

Behavioral sessions were recorded at 60 frames per second with OBS Studio software (version 27.1.3) with a USB camera (Arducam, Nanjing, China) positioned above the behavioral chamber. Videos were analyzed with DeepLabCut (version 2.2rc3) to track the 2-D position of mice during trials^13^, generating DeepLabCut-derived x/y positions and likelihood values for each video frame. These data were then processed using custom MATLAB code to find invalid points using the following three steps. First, any labeled points that had a likelihood value below 50% were removed. Second, labeled points that were positioned outside the bounds of the chamber (due to mislabeling the mouse’s reflection in the chamber wall) were removed. Finally, labeled points that moved >15 pixels in Euclidean distance between frames were removed. Invalidated points identified by these three steps were corrected using interpolation via piecewise cubit Hermite polynomials (*pchip*) to generate smoothed trajectories.

Mouse positional data (in pixels) were converted to real-world points (in millimeters) to allow comparisons between mice. Video calibration was achieved by recording videos of a checkerboard (1 x 1-cm squares) that was randomly moved within the behavioral chamber immediately before behavioral sessions began. Using randomly selected frames from these videos, built-in MATLAB functions (*detectCheckerboardPoints, estimateExtrinsics, img2world2d*) were used to estimate the camera position, to remove camera distortions, and to convert pixels to millimeters in DeepLabCut-generated trajectories. *Movement velocity* was calculated as the Euclidean distance in consecutive frames divided by the inter-frame interval (1/60 seconds).

### Stereotaxic surgery

Surgical methods were identical to those described previously^15,16^. Briefly, mice were anesthetized with inhaled 4% isoflurane and maintained at 1%–2% isoflurane during the surgery. Craniotomies were drilled above the prefrontal cortex (AP: +1.8, ML: ±0.4, DV: -1.7; 3 right hemisphere and 5 left hemisphere) or dorsomedial striatum (AP: +0.5, ML: ±1.4, DV: -2.7; 3 right hemisphere and 8 left hemisphere), and 4 × 4 electrode or optrode arrays were positioned at their target. Skull screws were used to anchor the surgical headcap and ground electrode arrays. Implants and screws were sealed with Slo-Zap cyanoacrylate (Pacer Technologies, Rancho Cucamonga, CA), accelerated with Zip-Kicker (Pacer Technologies), and methyl methacrylate (AM Systems, Port Angeles, WA). Mice were given at least 1 week for postoperative recovery before beginning food restriction, followed by training in the interval timing task.

### Neuronal recordings

Single neurons were recorded using a multi-electrode recording system (Open Ephys, Atlanta, GA). Recordings were processed using Plexon Offline Sorter (version 4.6.0, Plexon, Dallas, TX) to remove artifacts and single neurons were identified using PCA and waveform shape. The same neurons were sorted across interval timing sessions and Pavlovian conditioning sessions by loading data from both sessions together in Offline Sorter. Single neurons were identified based on the following criteria: 1) consistent waveform shape; 2) a separable cluster in PCA space; and 3) a consistent refractory period of at least 2 milliseconds in interspike interval histograms. Data for individual neurons were removed if the average firing rate was below 0.5 Hz over the entire behavioral session. Sorting integrity across the two types of behavioral sessions was quantified by comparing waveform similarity via correlation coefficients between the behavioral sessions. Data for neurons were removed if waveform correlation coefficient was below 0.8 (data for 14/291 neurons removed, all from DMS recordings). For recordings from DMS, putative medium spiny neurons were further separated from striatal fast-spiking interneurons based on hierarchical clustering of the waveform peak-to-trough ration and the half-peak width (*fitgmdist* and *cluster*)^4,5,16^. Only medium spiny neurons were analyzed from striatal recordings.

We calculated kernel density estimates of firing rates across the interval (-4 seconds before trial start to 4 seconds after trial end) binned at 0.2 seconds, with a bandwidth of 1. We used PCA to identify data-driven patterns of z-score neuronal prefrontal and striatal activity, as in our past work^4,5,7,15,16^. Based on this line of work, we focused on principal component 1 (PC1), which explained the most variance. PC1 exhibited time-dependent ramping activity, or a monotonic change across the temporal interval.

### Behavioral modulation patterns of neurons

To identify the modulation patters of individual neurons, we harnessed trial-by-trial generalized linear models (GLMS; *fitglme*) in MATLAB. For all GLMS, the response variable was firing rate; firing rate and all other events were binned at 0.2 seconds. For ramping neurons, we examined how a linearly increasing variable from 0 to 18 seconds over the interval predicted firing rate. For velocity modulation, we examined how velocity predicted firing rate. For nosepoke modulation, we examined how nosepoke entry predicted firing rate. For reward modulation, we examined how reward delivery predicted firing rate. For cue modulation, we examined how cue onset predicted firing rate. Only neurons with *p* values < 0.05 (tested using *anova*) after correction using Benjamini-Hochberg false discovery rate were considered to show significant modulation by time (i.e., ramping), velocity, nosepoke, reward, or cue. Slopes were derived for each neuron from the GLM fit of firing rate vs. time in the intervals over all trials. Adjusted R^2^ values were derived for each neuron from the GLM fit of firing rate vs. time (interval timing task or Pavlovian conditioning task) or velocity.

### Temporal decoding

To evaluate how neuronal ensembles predicted time, we constructed a naïve Bayesian classifier, following methods described previously^4,5,16,19^. Objective time was predicted from the firing rate during a trial using leave-one-out cross-validation. We ran the decoding analysis on different neuronal ensembles identified based on behavioral modulation patterns (i.e., ramping, velocity-modulated, nosepoke-modulated) or task (i.e., interval timing or Pavlovian conditioning). To correct for ensemble sizes, we ran the classifier on 1000 iterations of an equivalent number of randomly selected neurons from each group (20 neurons/group for ramping, velocity-modulated, and nosepoke-modulated neuronal ensembles, and 110 neurons/group for interval timing and Pavlovian conditioning ensembles) and quantified classifier performance by computing the R^2^ of objective time vs. predicted time for each trial. Classifier performance was compared to neuronal ensembles with time-shuffled firing rates.

### Immunohistochemistry

Mice were transcardially perfused with 1x phosphate-buffered saline (PBS) and 4% paraformaldehyde (PFA) after anesthesia using ketamine (100 mg/kg IP) and xylazine (10 mg/kg IP). Brains were fixed in a solution of 4% PFA followed by 30% sucrose in PBS before being cryosectioned. PFC-implanted brains were stained for tyrosine hydroxylase with primary antibody at 4 °C (rabbit anti-TH, Millipore AB152, 1:1000, 24 hours) and visualized with Alexa Fluor fluorescent secondary antibody at room temperature (goat anti-rabbit IgG Alexa 568, Invitrogen A11036, 1:1000, 2 hours). DMS-implanted brains were stained for tyrosine hydroxylase with primary antibodies at 4 °C (rabbit anti-TH, Millipore AB152, 1:1000, 24 hours or chicken anti-TH, Abcam ab76442, 1:1000, 48 hours) and visualized with Alexa Fluor fluorescent secondary antibodies at room temperature (goat anti-rabbit IgG Alexa 488, Invitrogen A11008, 1:1000, 2 hours or goat anti-chicken IgG Alexa 405, Invitrogen A48260, 1:250, 2 hours). To histologically confirm the location of electrode arrays, slices were imaged with VS-ASW-S6 imaging software (Olympus, Center Valley, PA).

### Statistics

We described data using group averages and standard error of the mean. To compare behavior between experimental groups, we used two-sample t-tests (*ttest2*) in MATLAB. To compare proportions of neuronal populations, we used Chi-squared tests (*crosstab*) in MATLAB. We compared PC1 scores and decoding accuracy (R^2^) by fitting linear models in R (*lmer*), running ANOVA to find main effects and/or interactions of neuron group and region, and making post hoc comparisons between groups using *emmeans*. Distance between adjusted R^2^ values were also compared between region by fitting linear models in R (*lmer*) and identifying main effects of region with ANOVA.

## RESULTS

### Prefrontal and striatal ensembles exhibit prominent time-related ramping

We recorded neuronal ensembles from the PFC or DMS of mice during interval timing task sessions^15,16,18^. In our mouse-optimized interval timing task, mice make a temporally-guided decision to switch their response from one nosepoke to another after they estimate that an interval of 6 seconds has elapsed (Fig 1A). Trials begin with a response at the back nosepoke which initiates tone and light cues. On 50% of randomly assigned trials, mice are rewarded for the first response at the short nosepoke after 6 seconds; these trials are not analyzed. On the other 50% of trials, mice must switch their response from the short nosepoke to the long nosepoke to be rewarded at the long nosepoke after 18 seconds. The switch *response time* is defined as the moment that mice exit the short nosepoke before responding at the long nosepoke. This response time reflects the mouse’s internal perception of elapsed time, because it is guided explicitly by temporal control of action rather than external cues. Accordingly, short nosepoke responses peaked at about 6 seconds and preceded switch responses, whereas long nosepoke responses peaked at closer to 18 seconds, when the reward became available (Fig 1B). There were no reliable behavioral differences between animals implanted with PFC or DMS recording electrodes (Fig S1).

We recorded 125 PFC neurons 152 DMS medium spiny neurons (MSNs) during interval timing (Fig S2A–B). As in previous work^2,4,20^, single PFC and DMS neurons displayed ramping activity, defined as a monotonic change in the firing rate across the 18-second interval (Fig 1C– F)^1,4,5,7,8^. Peri-event time histograms for PFC (Fig 1E) and DMS (Fig 1F) neurons showed a mixed distribution of neurons that ramped up vs. down in both populations. To characterize data-driven patterns of prefrontal and striatal activity, we used PCA^4,6^ to analyze PFC and DMS activity during the 18-second interval. As in our previous work^4,5,7,8,21^, principal component 1 (PC1) exhibited time-dependent ramping and explained the most variance (39%) in combined prefrontal and striatal activity (Fig 1G, Fig S2C–D). Consistent with past work, we found PC magnitudes were similar between PFC and DMS neurons (Fig 1H, PC1 |score|: t(275) = -0.7, *p* = 0.50; Fig S2E, PC2 |score|: t(275) = -0.8, *p* = 0.56)^4,5^.

To investigate single trial ramping dynamics, we turned to generalized linear models (GLMs, effect of time within the interval vs. trial-by-trial firing rate). Of note, PCA analysis identifies patterns that explain the most variance across prefrontal and striatal neuronal ensembles, whereas GLM analysis accounts for trial-by-trial patterns of activity and is more resistant to trial averaging^4,22^. Ramping neurons were defined by GLM fits and a false discovery rate (FDR)-corrected significant linear change in firing rate over the 18-second interval. We found that ramping activity was more frequent in DMS neurons (57%, 87/152 neurons) compared to PFC neurons (44%, 55/125 neurons; Fig 1I; χ^2^ = 4.8, *p* = 0.03). Accordingly, GLM-derived absolute linear slopes over time were greater for DMS neurons than for PFC neurons (Fig 1J; t(275) = -2.0, *p* = 0.04). As in prior work, these data indicate that ramping activity is prominent in the PFC and DMS during mouse-optimized interval timing task sessions^4,5,20^.

**Figure 1.**
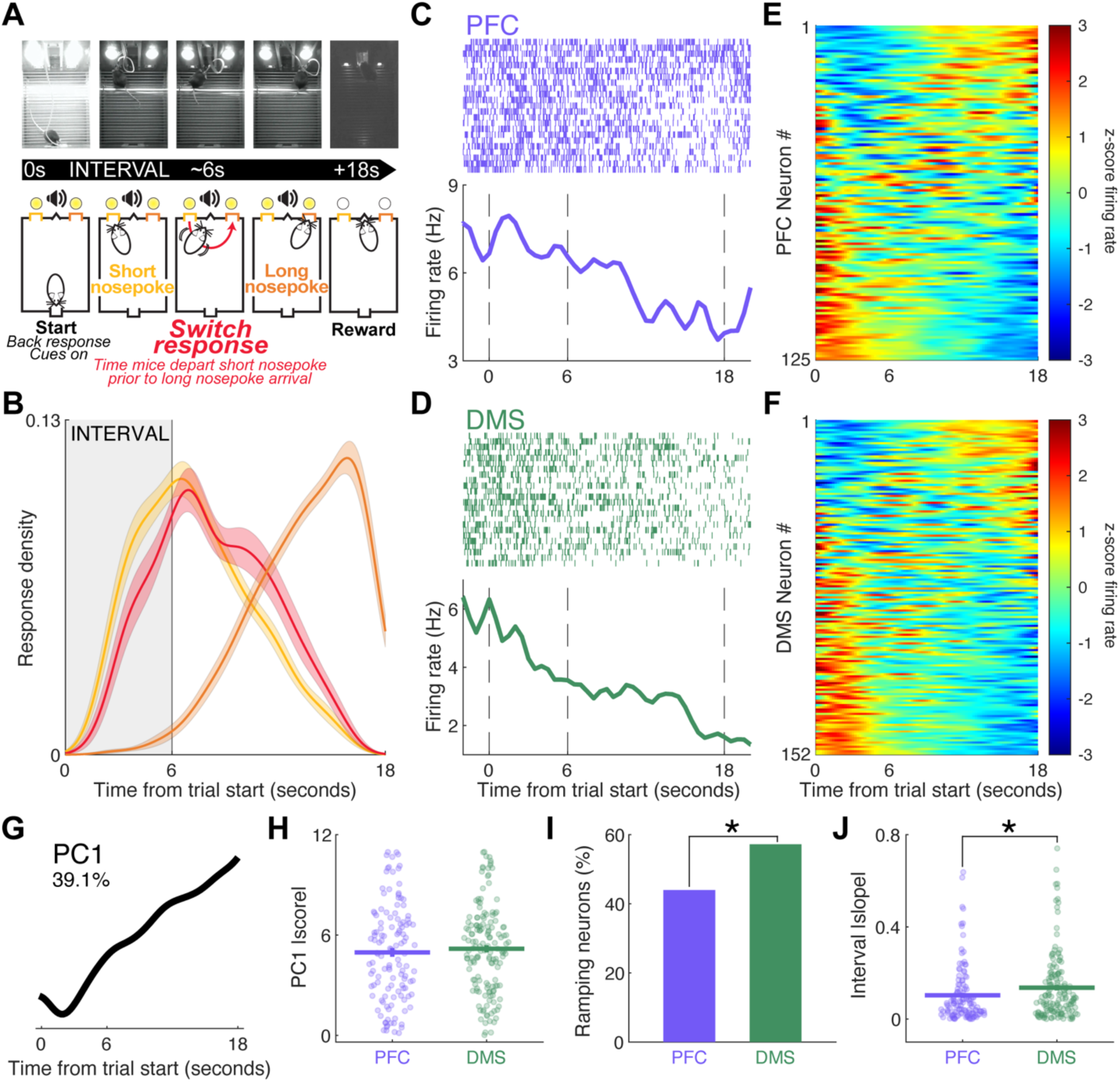
Mouse-optimized interval timing task. **A)** We trained mice to perform an interval timing task in which they switch from one nosepoke to another after estimating that a 6-second interval has elapsed. Mice initiate trials by responding at a back nosepoke. On 50% of trials, mice are rewarded for responding at the short nosepoke after 6 seconds; these trials are not analyzed. On the other 50% of trials, mice begin nosepoking at the short nosepoke but *respond* by switching to the long nosepoke after they estimate that 6 seconds has elapsed. Mice are rewarded for a response at the long nosepoke after 18 second. **B)** Response probability distribution from 19 mice (9 female) for short nosepokes (yellow), switch responses (red), and long nosepokes (orange). Shading represents standard error. **C)** Peri-event raster (top) and estimated average firing rate (bottom) of a time-modulated PFC neuron (purple) and **D)** a time-modulated DMS neuron (green). **E)** Peri-event time histograms (PETHs) from all PFC neurons and **F)** all DMS neurons. PETHs are sorted by principal component 1 (PC1) |score|. G) Principal component analysis (PCA) revealed that PC1 indicated time-dependent ramping activity and explained 39% of variance across combined PFC and DMS ensembles. H) PC1 |scores| did not differ between PFC (purple) and DMS (green) neurons. I) Ramping activity was observed in a greater proportion of neurons in the DMS compared to those in the PFC (\**p* < 0.05, x^2^). J) DMS neurons displayed a greater absolute linear slope (|slope|) than PFC neurons (\**p* < 0.05, t-test). In **H**) and **J**), dots represent individual neurons, horizontal lines represent group averages, and vertical lines represent standard error. Data from 125 PFC neurons in 8 mice and from 152 DMS neurons in 11 mice.

### Ramping neurons decode time more strongly than velocity-modulated neurons in striatum

Movement may contribute to prefrontal and striatal ramping activity^11^. To quantify motor contributions to ramping activity, we tracked the movement of mice during interval timing using DeepLabCut^13,14^. The movement trajectories of mice during interval timing task sessions were highly consistent (Fig2A–B). Movement velocity was greatest at ∼2 seconds, when mice moved from the back nosepoke to the short nosepoke (Fig 2C–D), and it did not differ between mice with PFC or DMS neuronal ensemble recordings (Fig 2E). Importantly, this movement velocity pattern was quite distinct from prefrontal and striatal neuronal activity quantified by PC1 (compare Fig 2E with Fig 1G).

To identify velocity-modulated neurons and ramping neurons, we used GLMs (effect of velocity or effect of time within the interval vs. trial-by-trial firing rate). We found that the proportion of velocity-modulated neurons and ramping neurons did not differ between mice with PFC or DMS neuronal ensemble recordings (Fig 2F-H; ramping neurons: χ^2^ = 2.1, *p* = 0.15; velocity-modulated neurons: χ^2^ = 1.3, *p* = 0.25; both: χ^2^ = 1.2, *p* = 0.28). However, the PFC contained significantly more velocity-modulated neurons than ramping neurons (Fig 2H; PFC: χ^2^ = 5.5, *p* = 0.02; DMS: χ^2^ = 0.02, *p* = 0.90). To verify ramping activity, we examined the PC1 |scores| of ramping vs. velocity-modulated neurons (Fig 2I). Critically, PC1 |scores| were higher for ramping neurons than for velocity-modulated neurons in both the PFC and the DMS (ramping vs. velocity: PFC: t(213) = 6.4, *p* < 0.0001; DMS: t(213) = 6.2, *p* < 0.0001), indicating that PC1 and GLMs capture similar ramping dynamics. Interestingly, neurons that were both ramping and velocity-modulated had lower PC1 scores in the DMS than those in the PFC (Fig 2I; PFC vs. DMS: t(213) = 3.1, *p* = 0.002). These data demonstrate that velocity modulations could not readily account for PC1 ramping activity.

To evaluate how well neuronal ensembles predicted time, we used a naïve Bayesian classifier, which generates trial-by-trial predictions of time from neuronal ensemble firing rates. To quantify temporal decoding, we computed the R^2^ of objective time vs. predicted time for each trial. Strikingly, DMS velocity-modulated neurons predicted time more weakly than ramping neurons (Fig 2J; ramping vs. velocity: t(228) = 8.7, *p* < 0.0001), while PFC temporal decoding was equivalent for ramping and velocity-modulated neurons (t(228) = 0.2, *p* = 1.00), and PFC ramping neurons decoded time poorly relative to DMS (ramping: t(228) = -2.8, *p* = 0.004; velocity: t(228) = 5.6, *p* < 0.0001). All ensembles predicted time significantly more strongly than their corresponding shuffled data (horizontal gray lines, *p* < 0.0001), suggesting some residual temporal decoding by velocity-modulated neurons. Together, these data suggest that striatal ramping and velocity modulations distinctly decode time, while PFC ramping and velocity modulations share temporal decoding.

**Figure 2.**
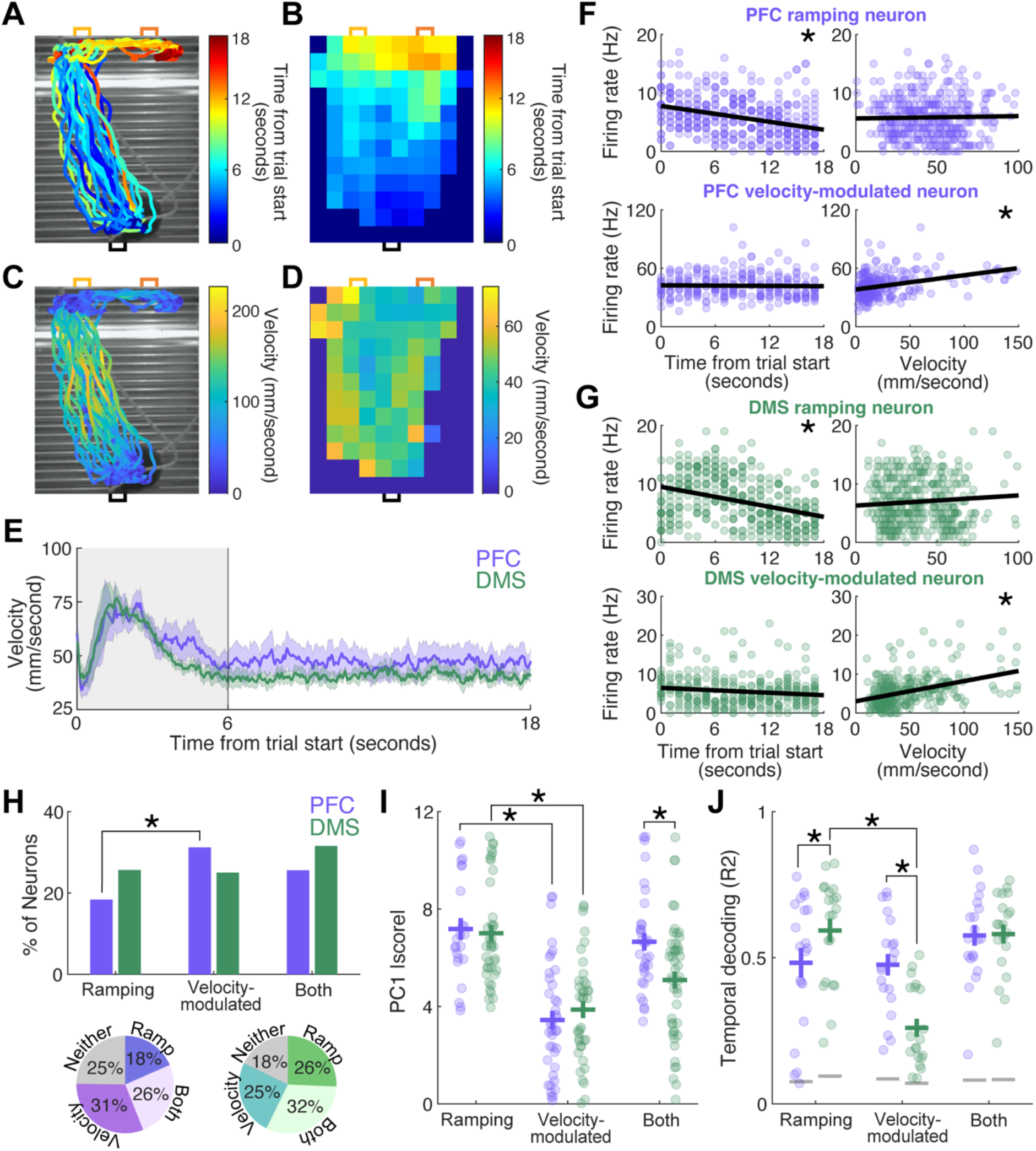
Ramping neurons decode time more strongly than velocity-modulated neurons. **A)** We tracked mouse position during trials using DeepLabCut^13^. Colored lines represent the movement trajectories of a single mouse during all interval timing switch trials from a single session. Color represents the time within the trial from trial start (seconds). **B)** Color represents the average time within the trial (seconds) when mice were within each block. **C)** We tracked mouse movement velocity using DeepLabCut. Colored lines represent the velocity of a single mouse during all interval timing switch trials from a single session. Color represents velocity (mm/second). **D)** Color represents the average movement velocity (mm/second) at which mice were moving within each part of the chamber. **E)** The average movement velocity (mm/second) during interval timing switch trials of mice with PFC (purple) vs. DMS (green) neuronal recordings. Shading represents s.e.m. **F)** PFC neurons could ramp without velocity modulation (top) or could have velocity modulation without ramping (bottom). Top row is the same neuron in left and right panels, and the bottom row is the same neuron in left and right panels. Dots represent firing rate and/or velocity binned into 1-second intervals. \**p* < 0.05 GLM. **G)** DMS neurons could ramp without velocity modulation (top) or could have velocity modulation without ramping (bottom). Top row is the same neuron in left and right panels, and the bottom row is the same neuron in left and right panels. Dots represent firing rate and/or velocity binned into 1-second intervals. \**p* < 0.05 GLM. **H)** Percentage of neurons in PFC (purple) and DMS (green) that are either ramping, modulated by velocity, or both. \**p* < 0.05, x^2^ PFC ramping vs. velocity-modulated neurons. Values in pie charts are rounded to the nearest whole number. **I)** PFC and DMS ramping neurons have stronger PC1 |scores| compared to PFC and DMS velocity-modulated neurons. The PC1 |scores| for neurons that are both ramping and velocity-modulated were lower in the DMS than in the PFC. \**p* < 0.05, two-way ANOVA with post hoc comparisons. Each dot represents an individual neuron, horizontal lines represent group means, vertical lines represent standard error. **J)** Ramping neurons in the DMS decoded time more strongly than those in the PFC and DMS ramping neurons decoded time more strongly than DMS velocity-modulated neurons. PFC velocity-modulated neurons decoded time more strongly than DMS velocity-modulated neurons. \**p* < 0.05, two-way ANOVA with post hoc comparisons. Each dot represents a trial, horizontal lines represent group means, and vertical lines represent standard error. Gray horizontal lines represent the mean for shuffled data. Data from 125 PFC neurons in 8 mice and 152 DMS neurons in 11 mice.

### Prefrontal and striatal ramping neurons decode time more strongly than nosepoke-modulated neurons

Other types of movements besides velocity could contribute to prefrontal and striatal ramping activity. We focused on nosepokes, the primary response-related movement that occurred during interval timing (Fig 3A–B, Fig S3A). We found that the percentage of nosepoke-modulated neurons did not differ between mice with PFC or DMS neuronal ensemble recordings (Fig 3C; nosepoke-modulated: χ^2^ = 0.1, *p* = 0.7; velocity-modulated: χ^2^ = 0.01, *p* = 0.91; both: χ^2^ = 0.09, *p* = 0.77). As with velocity-modulated neurons, the percentage of ramping and nosepoke-modulated neurons did not differ between PFC and DMS (Fig 3D; ramping: χ^2^ = 1.1, *p* = 0.30; nosepoke-modulated: χ^2^ = 2.1, *p* = 0.16; both: χ^2^ = 2.2, *p* = 0.14). Critically, as with velocity modulation, ramping activity as measured by PC1 |score| was significantly higher in ramping neurons than in nosepoke-modulated neurons (Fig 3E; ramping vs. nosepoke-modulated PFC: t(192) = 5.2, *p* < 0.0001; DMS: t(192) = 3.7, *p* = 0.001), verifying that PC1 and GLMs captured similar ramping dynamics. DMS neurons that were both ramping and nosepoke-modulated had lower PC1 |scores| than PFC neurons that were both ramping and nosepoke-modulated (PFC vs. DMS: both: t(192) = 2.14, *p* = 0.03).

Next, we examined temporal decoding for ramping and nosepoke-modulated neurons. We found that temporal decoding was significantly stronger for PFC and DMS ramping neurons than for nosepoke-modulated neurons (Fig 3F; PFC: t(228) = 3.8, *p* = 0.002; DMS: t(228) = 4.9, *p* < 0.0001). PFC neurons decoded time more strongly than DMS neurons in the ramping and nosepoke-modulated groups, whereas in the combined ramping and nosepoke-modulated neurons group, PFC neurons decoded time less strongly than DMS neurons (ramping: t(228) = 3.6, *p* = 0.0004; nosepoke-modulated: t(228) = 4.7, *p* < 0.0001; both: t(228) = -2.9, *p* = 0.004).

All neuronal ensemble groups predicted time significantly more strongly than their corresponding shuffled data (gray, *p* < 0.0001). When combined with comparisons of ramping and velocity-modulated neurons, these data provide further evidence that ramping and temporal encoding are distinct from movement-related activity in prefrontal and striatal ensembles.

**Figure 3.**
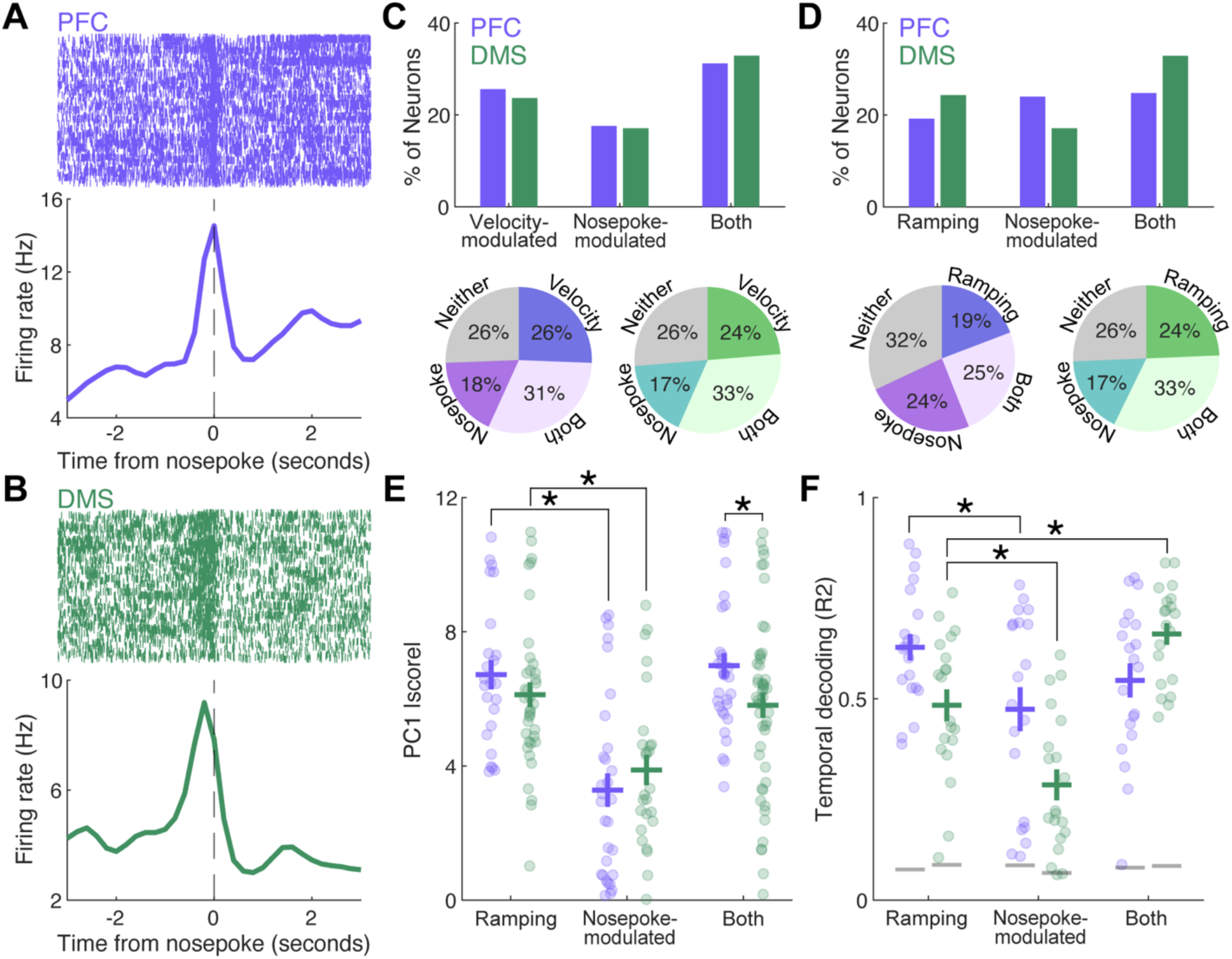
Ramping neurons decode time more strongly than nosepoke-modulated neurons. **A)** Top: Peri-event raster from a PFC neuron and **B)** a DMS neuron aligned to nosepoke responses across all interval timing trials from one session. Bottom: estimated mean firing rate. **C)** Percentage of neurons in PFC and DMS that are velocity-modulated, nosepoke-modulated, or both. Values in pie charts are ounded to the nearest whole number. **D)** Percentage of neurons in PFC and DMS that are ramping, nosepoke-modulated, or both. Values in pie charts are rounded to nearest whole number. **E)** PC1 |scores| for nosepoke-modulated neurons in the PFC and DMS are lower than for ramping neurons in the PFC and DMS. \**p* < 0.05, two-way ANOVA with post hoc comparisons. Each dot represents an individual neuron, horizontal lines represent group means, and vertical lines represent standard error. **F)** PFC and DMS ramping neurons decode time more strongly than PFC and DMS nosepoke-modulated neurons. DMS combined ramping and nosepoke-modulate neurons decode time more strongly than DMS ramping only neurons. \**p* < 0.05, two-way ANOVA with post hoc comparisons. Each dot represents a trial, horizontal lines represent group means, and vertical lines represent standard error. Gray horizontal lines represent means for shuffled data.

### Ramping activity is not associated with anticipation of reward delivery

Another potential contributor to ramping activity could be the anticipation of upcoming reward delivery. During the interval timing task, we found that 12% of PFC and 17% of DMS neurons were modulated by reward delivery (Fig S5B). To better isolate activity that anticipated reward delivery, we turned to Pavlovian conditioning, where a cue predicts reward delivery after a delay period^23^. We trained mice in the Pavlovian conditioning task after they had completed interval timing sessions. During Pavlovian conditioning, cues were randomly presented for 7 seconds, and mice were delivered a sucrose pellet reward immediately after the cues turned off (Fig 4A). Importantly, mice could not access nosepoke ports during Pavlovian conditioning sessions, and the auditory tone cue was changed to a different frequency to distinguish interval timing from Pavlovian conditioning sessions. Because mice were not required to make a nosepoke response to earn a reward, movement trajectories during Pavlovian conditioning trials were less consistent than during interval timing trials (Fig 4B compared to Fig 2A). PFC and DMS mice did not differ in their latency to retrieve reward following cues off (Fig 4C; t(17) = 1.0, *p* = 0.33).

We identified the same neurons that were recorded both during interval timing sessions and during Pavlovian conditioning sessions (Fig S4A–B; waveform correlations for PFC: R = 0.99 ± 0.002 and DMS: R = 0.99 ± 0.002). Most neurons had distinct activity between interval timing and Pavlovian conditioning (Fig 4D–G). Comparing activity from interval timing and the Pavlovian conditioning task demonstrated that ramping was rare during the Pavlovian conditioning task (Fig 4E, G), as there were fewer ramping neurons during Pavlovian conditioning compared to interval timing (Fig 4H; PFC: χ^2^ = 25.4, *p* = 4.70x10^-7^; DMS: χ^2^ = 27.9, *p* = 1.30x10^-7^). Only 6% of PFC neurons and 14% of DMS neurons ramped during both tasks, which was less than during interval timing alone (Timing vs. Both: PFC: χ^2^ = 31.5, *p* = 2.03x10^-8^; DMS: χ^2^ = 27.9, *p* = 1.30x10^-7^). By comparison, the proportion of neurons that responded to cues at trial start did not differ between tasks (Fig S5A; PFC: χ^2^ = 1.2, *p* = 0.265; DMS: χ^2^ = 0.36, *p* = 0.548), although only 6% of PFC neurons and 7% of DMS neurons were cue-responsive across both tasks (Timing vs. Both: PFC: χ^2^ = 6.6, *p* = 0.01; DMS: χ^2^ = 6.18, *p* = 0.013). Similarly, more neurons were reward-responsive during interval timing than during Pavlovian conditioning (Fig S5B; PFC: χ^2^ = 6.9, *p* = 8.66 x 10^-3^; DMS: χ^2^ = 9.3, *p* = 2.25 x 10^-3^), and no neurons were reward-responsive across both tasks. Notably, we found that ramping activity was largely stable between two distinct interval timing sessions on separate days (Fig S4C–E). These data demonstrate that temporal modulations during Pavlovian conditioning are sparse, while other modulations are preserved.

Finally, we compared temporal decoding occurring during interval timing trials vs. during Pavlovian conditioning trials. Temporal decoding was significantly weaker during Pavlovian conditioning compared to interval timing for both the PFC and the DMS (Fig 4I; 0–18 seconds: PFC: t(228) = 8.1, *p* < 0.0001; DMS: t(228) = 5.3, *p* < 0.0001; 0–6 seconds: PFC: t(228) = 5.6, *p* <0.0001; DMS: t(228) = 5.0, *p* < 0.0001). During Pavlovian conditioning trials, DMS ensembles decoded time more strongly than PFC ensembles (t(228) = -2.5, *p* = 0.01). All recorded ensembles predicted time significantly more strongly than their corresponding shuffled data (gray bars). These results suggest that PFC and DMS ensembles encode temporal intervals only when the mouse must cognitively control their actions in time to receive reward. These data show that ramping activity is distinct from reward-related activity in prefrontal and striatal ensembles.

**Figure 4.**
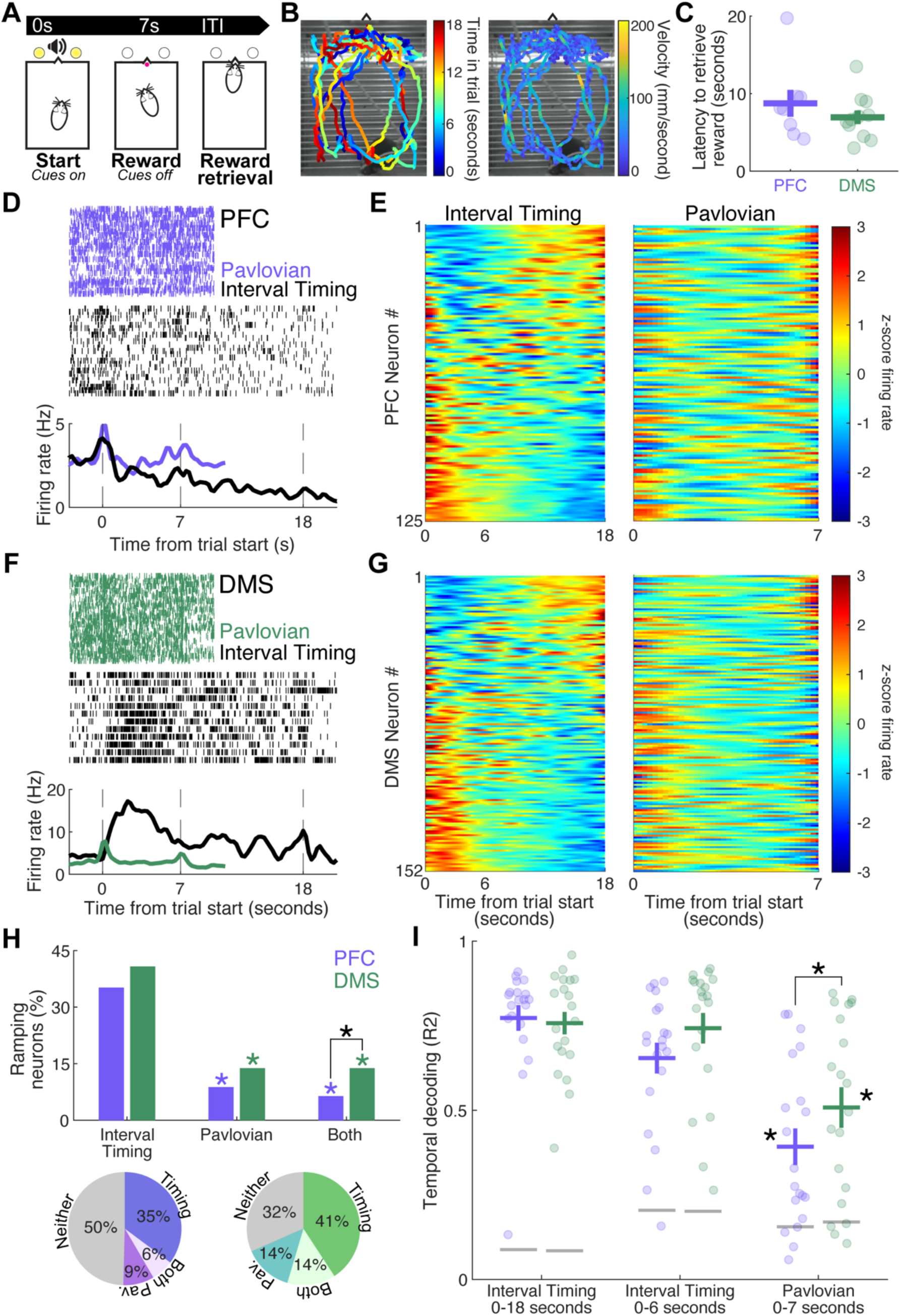
Ramping activity is not associated with anticipation of reward delivery. **A)** Following recordings in the interval timing task, we trained mice in a Pavlovian conditioning task. In this task, cues turn on for 7 seconds, and a sucrose pellet is delivered after cues are turned off. **B)** Example movement trajectories from two mice during Pavlovian conditioning trials. Color represents the time within the trial (seconds). **C)** PFC and DMS mice did not differ in their latency to retrieve reward. Each dot represents the average latency of an individual mouse, horizontal lines represent group means, and vertical lines represent standard error. **D)** Top: Peri-event raster of Pavlovian conditioning trials (purple) and interval timing switch trials (black) for the same PFC neuron. Bottom: Estimated mean firing rate across all trials for both tasks. **E)** Left: Peri-event time histogram (PETH) of switch trials during interval timing sessions for all PFC neurons, sorted by PC1. Right: PETH of Pavlovian conditioning trials for all PFC neurons. Neurons are displayed in the same order as the neurons in the interval timing PETH. **F)** Top: Peri-event raster of Pavlovian conditioning trials (green) and interval timing switch trials (black) for the same DMS neuron. Bottom: Estimated mean firing rate across all trials for both tasks. **G)** Left: PETH of switch trials during interval timing sessions for all DMS neurons, sorted by PC1. Right: PETH of Pavlovian conditioning trials for all DMS neurons. Neurons are displayed in the same order as the neurons in the interval timing PETH. **H)** Percentage of PFC (purple) and DMS (green) ramping neurons during the interval timing task (0–18 seconds), the Pavlovian conditioning task (0–7 seconds), both tasks, or that did not ramp during either task (neither). Values in pie charts are rounded to the nearest whole number. **I)** Temporal decoding was stronger in PFC and DMS ensembles during interval timing trials (0– 18 and 0–6 seconds) than in Pavlovian conditioning trials (0–7 seconds). DMS ensembles had stronger temporal decoding during Pavlovian conditioning than PFC ensembles. Temporal decoding was stronger from both PFC and DMS ensembles in non-shuffled task data than in shuffled data, \**p* < 0.05, two-way ANOVA with post hoc contrast testing. Each dot represents a single trial of behavior for PFC ensembles (8 mice) and DMS ensembles (11 mice). Horizontal lines represent mean, and vertical lines represent standard error.

### Timing, movement, and reward contributions to prefrontal and striatal activity

Our work suggests that timing, movement, and reward signals exist within the PFC and DMS. To quantify the relative contribution of these factors, we calculated the adjusted R^2^ of ramping, velocity, and reward anticipation to the firing rate of each recorded PFC and DMS neuron (Fig 5A–B). DMS neurons (Fig 5B) appeared to more broadly represent different behavioral components than PFC neurons (Fig 5A). This was driven primarily by the contribution of reward anticipation to the DMS, as the Euclidean distance between the R^2^ values for ramping vs. Pavlovian conditioning was significantly greater for DMS neurons than for PFC neurons (Fig 5C; F(1) = 4.6, *p* = 0.03), but it was not different between the R^2^ values for ramping vs. velocity (Fig 5D; F(1) = 1.5, *p* = 0.28). Taken together, our results quantify behavioral contributions to prefrontal and striatal ensembles.

**Figure 5.**
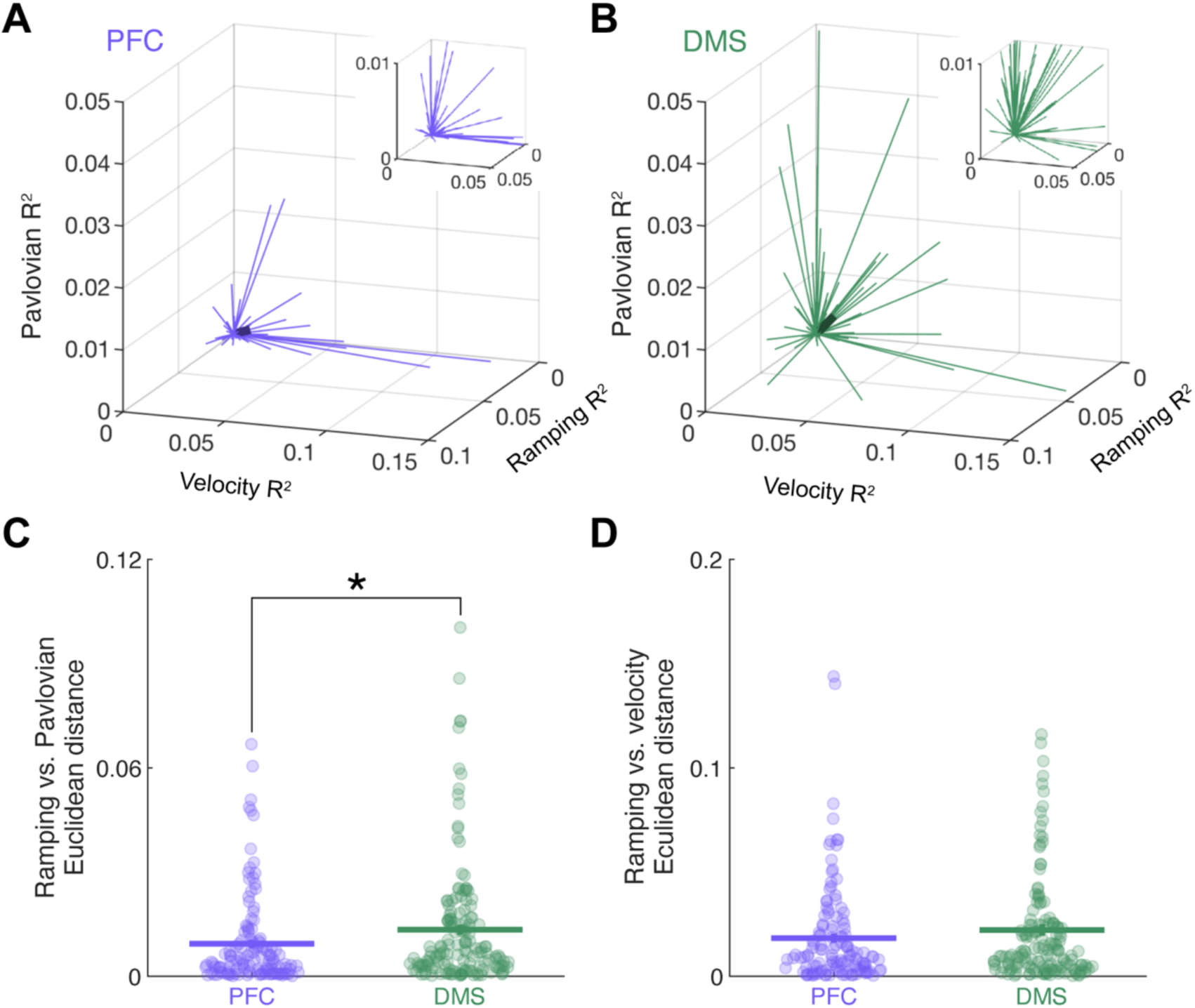
Timing, movement, and reward contributions to prefrontal and striatal activity. **A)** Explained variance (adjusted R^2^) from ramping, velocity, and Pavlovian conditioning (reward anticipation) for each recorded PFC neuron. Each thin purple line represents a single neuron. The thick, dark purple line represents the average. Inset: magnified area around the origin to show neurons with smaller adjusted R^2^ values. Data from 125 neurons (8 mice). **B)** Explained variance (adjusted R^2^) from ramping, velocity, and Pavlovian conditioning (reward anticipation) for each recorded DMS neuron. Each thin green line represents a single neuron. The thick, dark green line represents the average. Inset: magnified area around the origin to show neurons with smaller adjusted R^2^ values. Data from 152 neurons (11 mice). **C)** The Euclidean distance between R^2^ values for ramping vs. Pavlovian conditioning was significantly greater in DMS neurons than in PFC neurons. \**p* < 0.0.5 ANOVA. Each dot represents a single neuron, horizontal lines represent group averages, and vertical lines represent standard error. **D)** The Euclidean distance between R^2^ values for ramping vs. velocity did not differ between PFC and DMS neurons. Each dot represents a single neuron, horizontal lines represent group averages, and vertical lines represent standard error.

## DISCUSSION

Ramping is a prominent component of prefrontal and striatal neuronal activity during motivated behavior^1^. It is observed across species when participants must guide their actions in time^2,4,5,20^, and explicitly correlates with internal temporal estimates^2,5^. Ramping activity is also observed during other cognitive operations, such as working memory or reversal learning^24–26^. Despite its prominence, it is unclear what specific behaviors contribute to ramping activity. We quantified the contribution of movement (velocity), task-specific movement<u>s</u> (nosepoke), and reward anticipation to ramping activity in PFC and DMS neurons using two novel methods. First, we used DeepLabCut to track movement of mice during interval timing trials and found stronger temporal decoding associated with ramping activity than with movement-related activity, with the exception of velocity-modulated neurons in the PFC. Second, we trained animals in both an interval timing task and a Pavlovian conditioning task and found stronger ramping during interval timing trials than during Pavlovian conditioning trials. Our results demonstrate that movement and reward-anticipation could not fully account for temporal decoding of ramping neurons in the PFC or DMS, suggesting that ramping is a critical temporal signal in prefrontal and striatal neuronal activity. Together, our data indicate that prefrontal and striatal ramping is a cognitive signal that actively estimates time rather than a movement or reward-anticipation signal.

Ramping activity can be captured by drift-diffusion computational models, in which activity “drifts” from a starting point toward a decision threshold^9,16,27^. Ramping dynamics can be integrated and compared to a decision threshold^28^. Decision thresholds can be modulated by brain areas downstream from corticostriatal neuronal ensembles, such as the subthalamic nucleus^29,30^. Our data support drift-diffusion models of temporal encoding: in both the PFC and DMS, temporal decoding was stronger for ramping neurons than for movement-or reward-modulated neurons, with the exception of velocity-modulated neurons in the PFC. Similarly, in our prior work, temporal decoding was weaker in ensembles of non-ramping neurons compared to ramping neurons^5,31^. Furthermore, our prior work found that manipulations that decrease corticostriatal ramping also decrease temporal decoding^5,8,16^. Together, these data support the idea that prefrontal and striatal neuronal ensembles represent time through drift-diffusion dynamics.

An alternative to the drift-diffusion model is that neuronal ensembles represent time by a distributed population code—that is, at each moment, a neuronal ensemble is in a distinct state, and neuronal decoders can readily read out time from these ensemble states^19,20,32,33^. We found that during interval timing, ensembles comprising velocity-modulated and nosepoke-modulated neurons outperformed shuffled data, as did neuronal ensembles during Pavlovian conditioning trials. These findings suggest that residual temporal decoding exists without ramping, providing some support for population coding. While prefrontal and striatal ensembles generally showed much stronger temporal decoding associated with ramping activity than with movement or reward modulation, it is unclear whether mammalian brains harness drift-diffusion dynamics, distributed population codes, both, or other temporal encoding schemes. Resolving these possibilities will require future experiments that specifically isolate and perturb these models.

Our findings quantify the contribution of coarse movements such as velocity and nosepokes to PFC and DMS ramping. It has been proposed that temporal signals are embodied in movements^11,34^. However, three lines of evidence argue that gross movements do not entirely explain ramping activity. 1) We find that ramping activity decodes time much more strongly than movement-related signals. 2) In prior studies, ramping activity was observed even during “hold” periods when animals are not moving, including head-fixed behavior^7,35,36^. 3) We found that velocity or nosepokes were quite distinct from ramping or response times. Although nosepoking and moving around the chamber are two of the most prominent mouse movement<u>s</u>, we did not track fine movements which might track or reflect time. Future studies are needed to more precisely measure movements or to pursue further behavioral control in animals that are required to suppress movements to time.

Dopamine and neuronal activity can ramp toward reward^12,37,38^. Another novel aspect of our study is that it compared activity in the same neurons and animals that performed tasks in two different contexts: operant-based interval timing, where mice must actively track time to earn a reward, and classical Pavlovian conditioning, where mice do not need to track time but learn to anticipate when reward will be delivered. Importantly, during interval timing mice must actively track time to determine when to switch from the short to the long nosepoke, whereas during the Pavlovian conditioning task, rewards are always delivered after 7 seconds, and mice may be less engaged. While corticostriatal networks can represent the temporal statistics of upcoming rewards^39^, our data suggest that ramping activity and temporal decoding are much stronger during operant-based interval timing. This further argues that prefrontal and striatal ramping activity is a signature of cognitive activity, because ramping was significantly diminished when mice were not required to actively track time.

Our work has three limitations. First, our recent work demonstrated that molecular and anatomical factors, which we did not explore here, further explain temporal patterns within corticostriatal circuits^16,40^. However, identifying these subpopulations requires optogenetic tagging or cell type-specific fiber photometry. Second, we did not track fine movements, and we did not use behavioral tasks that formally control for movements, such as temporal bisection tasks^33^. However, rodent peak interval timing directly translates to human diseases such as schizophrenia and Parkinson’s disease^5,8,41–43^, making it particularly important to establish motor and reward contributions to neuronal activity during this version of interval timing. Finally, we did not systematically vary reward valence or explore negative reinforcers, which may change the salience of Pavlovian conditioning and vary temporal expectation.

In summary, we quantified the contribution of time, movement, and reward anticipation to ramping activity in prefrontal and striatal neuronal ensembles. We compared neuronal activity in the PFC and DMS of mice performing an interval timing task (with motion tracking) and in the same mice performing a Pavlovian conditioning task. We found that ramping activity decodes time significantly more strongly than movement-related or reward-related signals.

These data help us understand and quantify the contributions of time, movement, and reward to corticostriatal neuronal activity during temporal control of action, and they suggest that ramping activity represents time rather than movement or reward.

## SUPPLEMENTARY FIGURES

**Figure S1.**
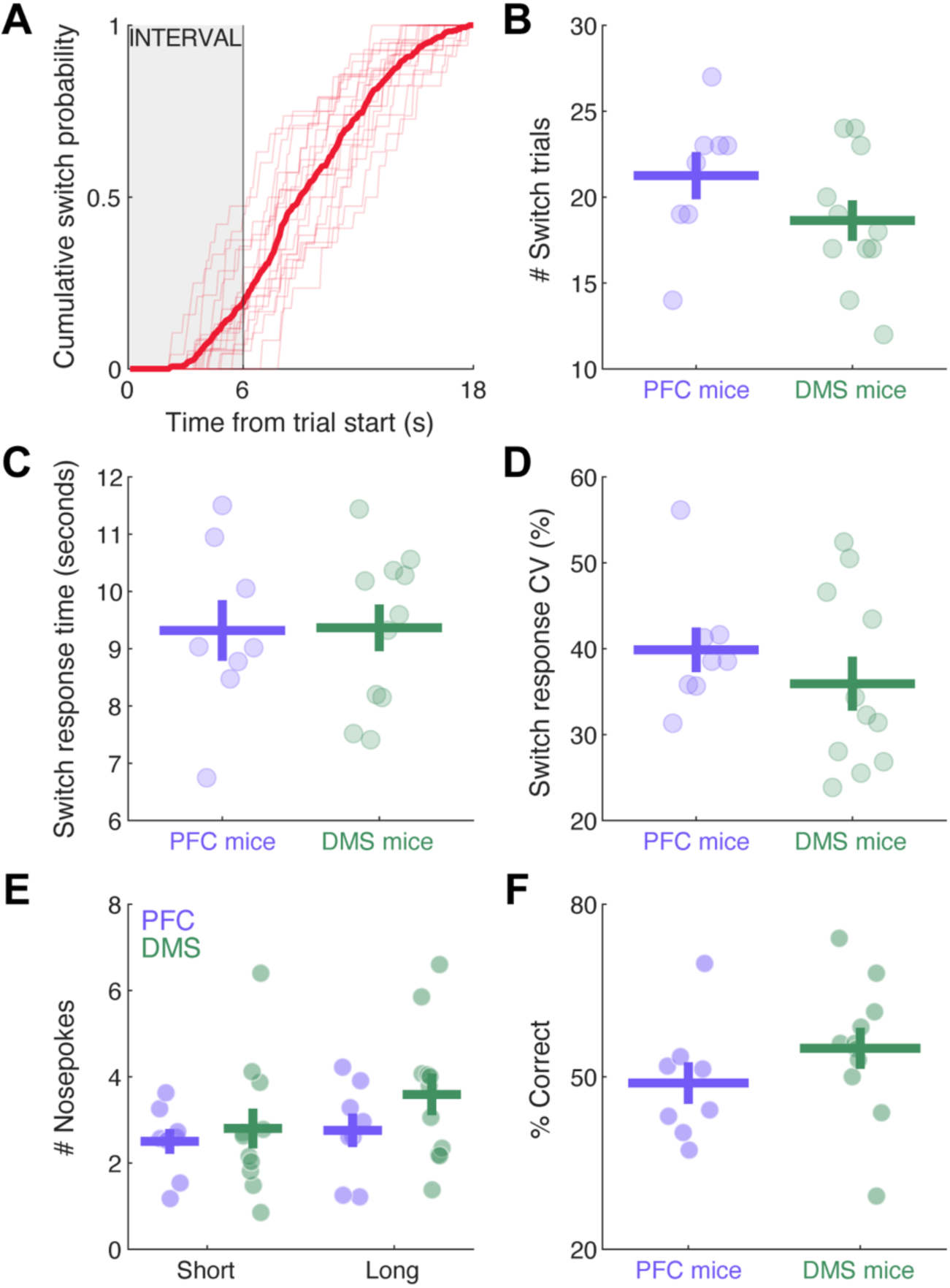
Interval timing behavior. **A)** Cumulative switch response density of 19 mice (8 PFC and 11 DMS). The thin lines represent individual mice, and the thick line represents the average. **B)** PFC mice and DMS mice performed a similar number of switch trials per session (t(17) = 1.4, *p* = 0.17). **C)** PFC mice and DMS mice had similar average switch response times (t(17) = -0.1, *p* = 0.95). **D)** PFC mice and DMS mice had similar switch response coefficients of variation (CV; t(17) = 0.9, *p* = 0.38). **E)** PFC mice and DMS mice made a similar number of nosepokes during switch trials (Short: t(17) = -0.5, *p* = 0.62; Long: t(17) = -1.3, *p* = 0.22). **F)** PFC mice and DMS mice correctly performed a similar percentage of switch trials (t(17) = - 1.2, *p* = 0.26). These data suggest that interval timing performance was highly consistent across mice from these two groups^16,17,31,44,45^.

**Figure S2.**
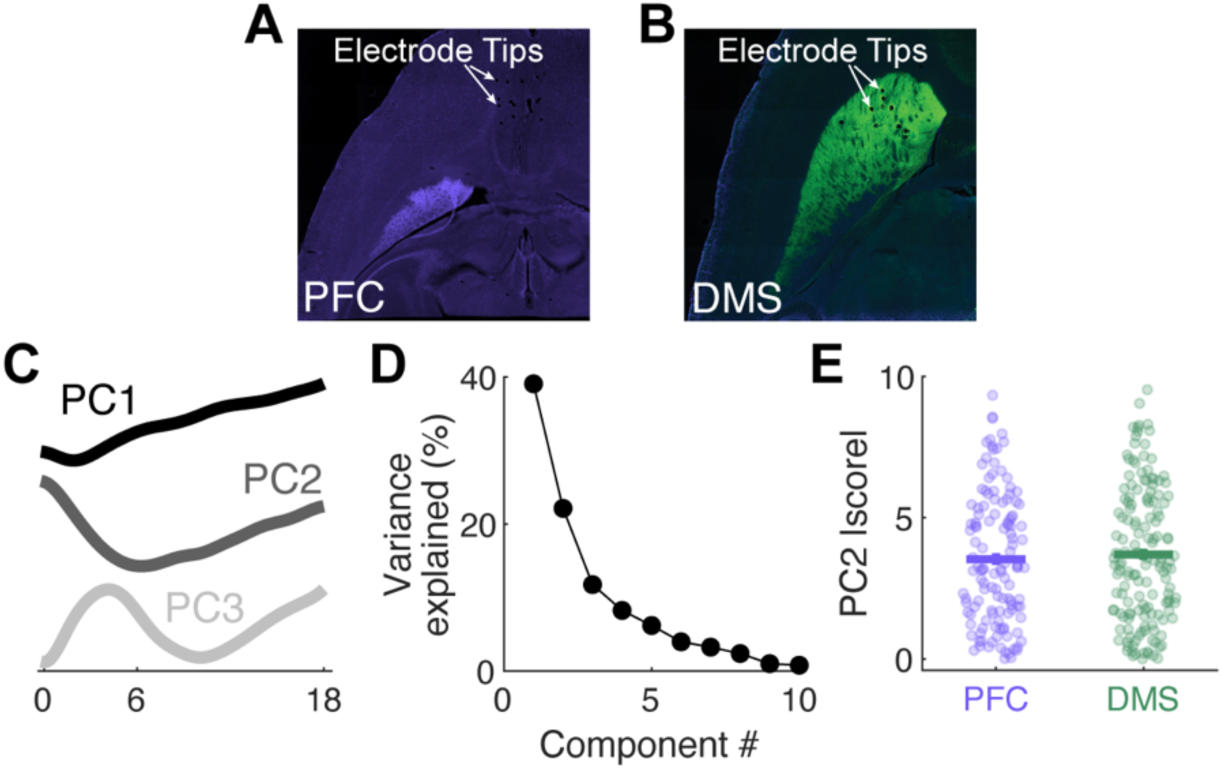
PFC and DMS principal components. **A)** Prefrontal (PFC) recording array locations. **B)** Dorsomedial striatum (DMS) recording locations. **C)** Principal component 1 (PC1)–PC3 of PFC and DMS neurons together. **D)** Variance explained by PC1–PC10. **E)** PC2 did not differ between PFC and DMS neurons (t(275) = -5.9, *p* = 0.56).

**Figure S3.**
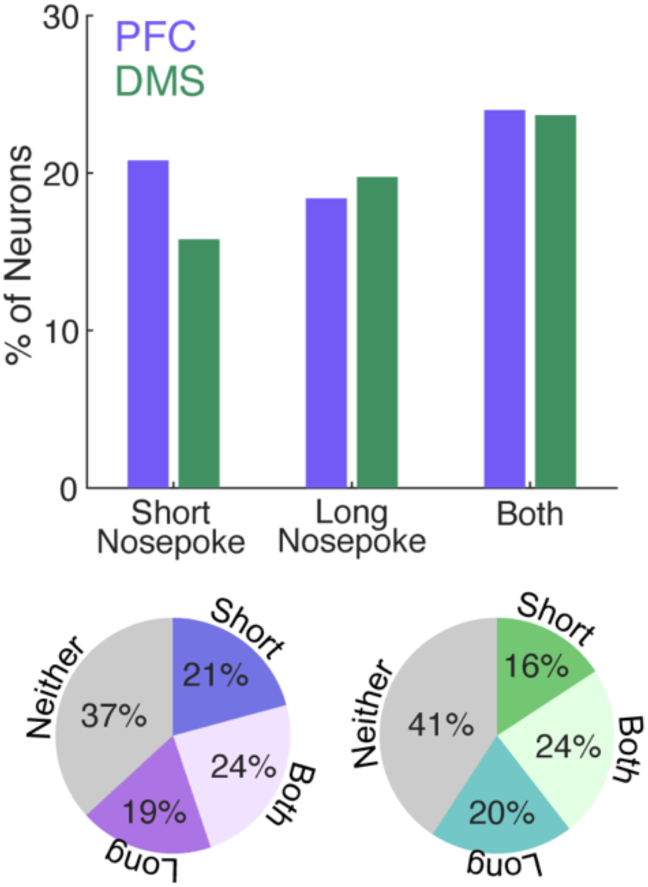
PFC and DMS nosepoke modulation. Percentage of PFC (purple) and DMS (green) neurons that displayed modulation, grouped by short nosepokes, long nosepokes, both short and long nosepokes, or neither.

**Figure S4.**
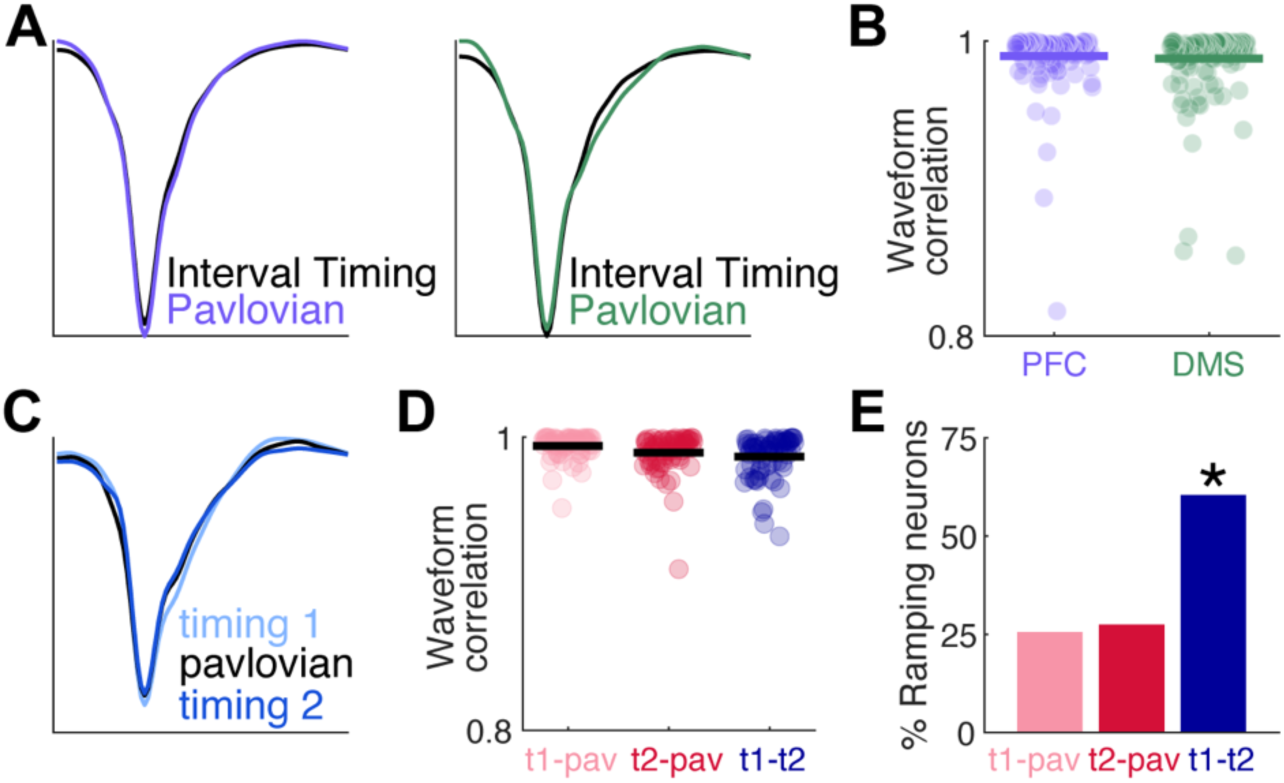
Ramping is stable within neurons across interval timing sessions. **A)** Example waveforms averaged across all action potentials from one PFC neuron (left) and one DMS neuron (right) in an interval timing session (black) and a Pavlovian conditioning session (purple (left) or green (right)). **B)** Waveforms were highly correlated between interval timing and Pavlovian conditioning sessions. Each dot represents a single neuron, and the horizontal lines represent the mean. **C)** Example waveform of a neuron recorded across three separate behavior sessions: interval timing session 1 (light blue), Pavlovian conditioning session (black), and interval timing session 2 (dark blue). **D)** Waveforms were highly correlated across all sessions. Each dot represents a single neuron, and the horizontal lines represent average correlations. **E)** While only ∼25% of neurons displayed ramping activity in both interval timing and Pavlovian conditioning sessions, ∼60% of neurons maintained ramping activity between the two interval timing sessions (t1-pav vs. t1-t2: χ^2^ = 1.1, *p* = 9.93x10^-4^; t2-pav vs. t1-t2: χ^2^ = 1.3, *p* = 4.17x10^-4^).

**Figure S5.**
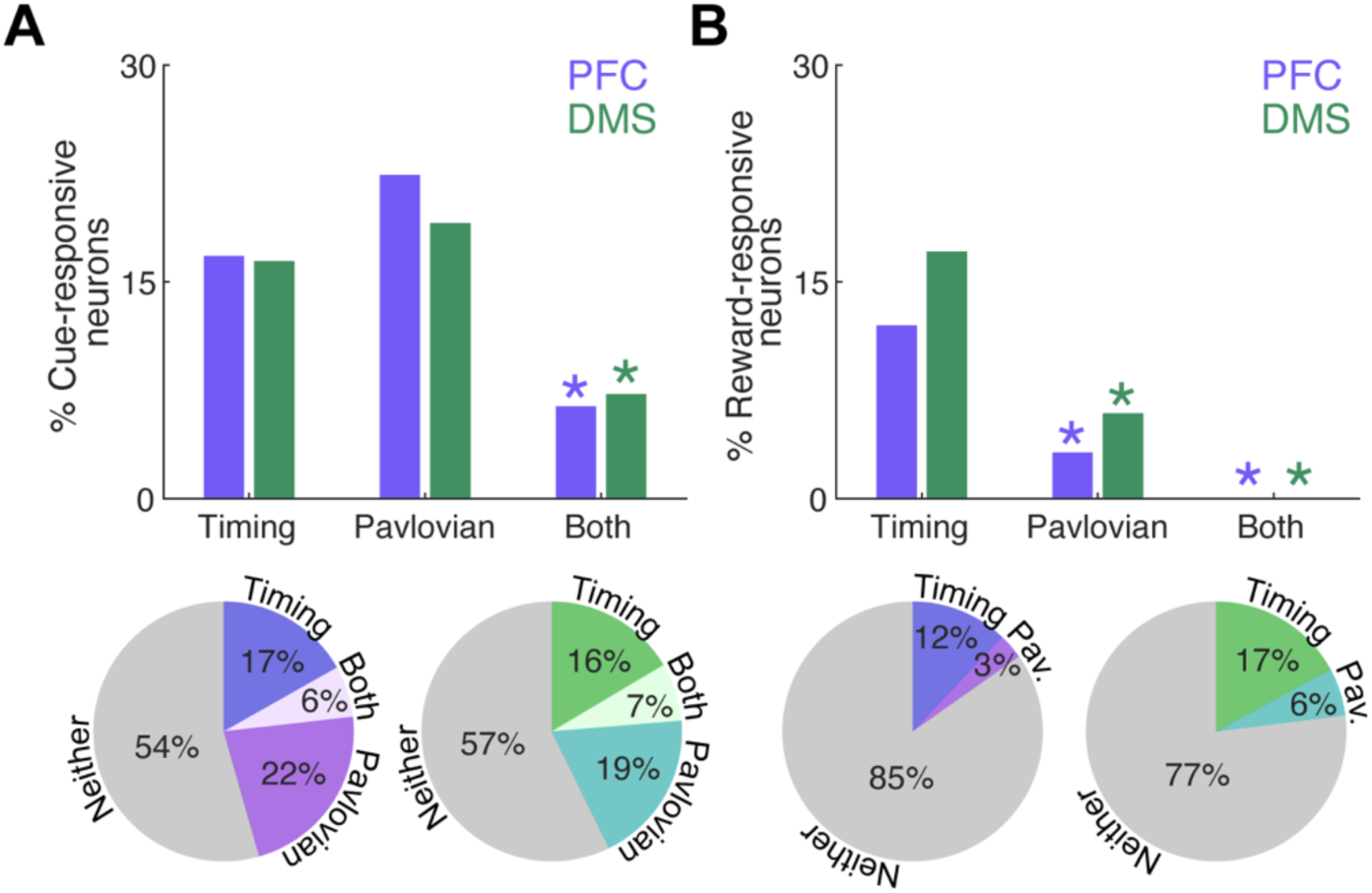
PFC and DMS cue and reward modulation. **A)** A similar proportion of neurons responded to cue onset during interval timing and Pavlovian conditioning tasks in the PFC (purple) and DMS (green). Significantly fewer neurons were responsive to cues in both types of tasks than in interval timing only. \**p* < 0.05, χ^2^ **B)** Significantly fewer neurons were modulated by reward delivery during the Pavlovian conditioning task than during the interval timing task. No neurons were reward-responsive during both types of tasks. *p < 0.05, χ^2^

